# Novel haplotypes and networks of *AVR-Pik* alleles in *Magnaporthe oryzae*

**DOI:** 10.1101/499335

**Authors:** Jinbin Li, Qun Wang, Chengyun Li, Yunqing Bi, Xue Fu, Raoquan Wang

## Abstract

Li J, Wang Q, Li C, Bi Y, Fu X, Wang R. 2018. Novel haplotypes and networks of *AVR-Pik* alleles in *Magnaporthe oryzae*. PLoS Pathog _______.

Rice blast disease is one of the most destructive fungal diseases of rice world-wide. The avirulence (*AVR*) genes of *Magnaporthe oryzae* are recognized by the cognate resistance (*R*) genes of rice, and trigger race specific resistance. Here, we studied the possible evolutionary pathways in the evolution of *AVR-Pik* alleles by analyzing the DNA sequence variation and assayed for their avirulence function to the cognate *Pik* alleles resistance genes under field conditions in China. Results of PCR products showed that 278 isolates of *M. oryzae* carry *AVR-Pik* alleles among genomic DNA of 366 isolates of *M. oryzae* collected from Yunnan Province, China. Among of them, 66.7-90.3% of *M. oryzae* carry *AVR-Pik* alleles from six regions of Yunnan. Moreover, 10 *AVR-Pik* haplotypes encoding five novel *AVR-Pik* variants were identified among 201 isolates. The *AVR-Pik* alleles stepwise evolved to virulence from avirulent forms via base substitution. These findings demonstrate that *AVR-Pik* alleles are under positive selection and mutations are responsible for defeating race-specific resistance *Pik* alleles in nature.

**Author summary:** The interaction of resistance gene (R) of rice and avirulence (*AVR*) gene of rice blast fungi are belong to the gene-for-gene theory, the variation of *AVR* is one of the major reasons for generation new race. To detect the variation of *AVR* gene in isolates population of *Magnaporthe oryzae* collected from rice production fields, will helpful for evaluated the effectiveness of *R* gens in rice production areas. The *Pik* allele contained five *R* genes of *Pik, Pikh, Pikp, Pikm* and *Piks*, and the corresponding to the *AVR* genes of *AVR-Pik/kh/kp/km/ks* of *M. oryzae*. The *Pik* gene specifically recognizes and prevents infections by isolates of *M. oryzae* that contain *AVR-Pik*. The molecular variation of *AVR-Pik* alleles of *M. oryzae* and *Pik* alleles of rice remains unclear. Here we demonstrated that polymorphism and distribution of *AVR-Pik* alleles in Yunnan Province, China. By pathogenicity assays to detect function of the different haplotypes of *AVR*-*Pik*, for the first time we show detour and stepwise evolution of *AVR-Pik* alleles in rice production areas of Yunnan. The functional *AVR-Pik* possesses diversified sequence structures, and is under positive selection pressure in nature.

## Introduction

In the long history of coexistence of parasitism and predation, the adaptive genetic mutation are including between hosts and pathogen. While the selective pressure was considered as the main force. So far, two hypotheses of arms race and trench warfare evolution have been proposed between hosts resistance genes (*R*) and pathogens avirulence genes (*AVR*).[1] The arms race was received as principal hypothesis, in which both hosts *R* and pathogens *AVR* were under directional selection, and the alleles derived by mutation, in brief, pathogen evolved a virulence gene in order to overcome the host defense, on the other hand, the hosts evolved a new resistance allele to defeat the virulence genes of the pathogen. Where as, in trench warfare hypothesis, the evolution of both hosts *R* and pathogens *AVR* is non-directional.

Rice blast is one of the most destructive diseases of rice growing regions, caused by the filamentous ascomycetous fungi *Magnaporthe oryzae* (synonym of *Pyricularia oryzae*). Employing resistant rice varieties with major resistance (R) gene was considered of the most important strategy to control this disease, which is environmentally friendly and economical for decreasing crop loss by rice blast. So far, ≤26 *R* genes in rice have been cloned: *Pb1, Pia, Pib, Pid2, Pid3, Pik, Pikh/Pi54, Pikm, Pikp, Pish, Pit, Pita, Pizt, Pi1, Pi2, Pi5, Pi9, pi21, Pi25, Pi36, Pi37, Pi56, Pi63, PiCO39* (http://www.ricedata.cn/gene/gene_pi.htm), *Pi64*[2] and Pigm[3].

Rice resistance gene can recognize the corresponding *AVR* of *M. oryzae* and triggers the immunity reaction. So far, 12 *AVR* genes in *M. oryzae* have been cloned: *AVR-Pi54*[4], *AVR-Pi9*[5], *AVR-Pib*[6], *AVR-Pia*[7], *AVR-Pii*[7], *AVR-Pik/km/kp*[7], *AVR-Pizt*[8], *ACEl*[9], *AVR-Pita*[10], *AVR1-CO39*[11], *PWL1*[12], and *PWL2*[13]. The *AVR-Pik/km/kp* gene of *M. oryzae* inspects the effectiveness of *R* gene *Pik/km/kp. AVR-Pik/km/kp* encodes a putative secreted protein with 113 amino acids containing two conserved motifs: motif-1, [LI]xAR[SE][DSE]; and motif-2, [RK]CxxCxxxxxxxxxxxxH (similarity to the C2H2 zinc finger motif). [7] The *AVR-Pik/km/kp* gene was cloned from the isolate of Ina168 but absent in the assembled sequence of isolate 70-15, which is recognized by host resistance of Pik protein and triggers the defense response.[7] Five *AVR-Pik* alleles (*AVR-Pik-A, AVR-Pik-B, AVR-Pik-C, AVR-Pik-D, AVR-Pik-E*) were found[7], and the *AVR-Pik-D* (20.5%) and *AVR-Pik-E* (1.4%) was detected among 77 isolates.[14] Four *AVR-Pik* alleles (*AVR-Pik-A, AVR-Pik-C, AVR-Pik-D, AVR-Pik-E*)were found among 39 isolates from world-wide (three isolates from Europe, six isolates from America, seven isolates from Africa and 23 isolates from Asia), and the *AVR-Pik-D* was the highest frequent allele (15 out 39), while the *AVR-Pik-A, AVR-Pik-C, AVR-Pik-E* alleles had similar frequencies (7-9 out of 39).[15] *AVR-Pik/km/kp* has evolved via gene gain/loss manners[7], while substitution mutations were observed in coding regions of the *AVR-Pik/km/kp* in *M. oryzae* populations and 16 SNPs were found in non-signal domain harbored regions in Chinese rice blast isolates.[16]

Rice resistance gene can recognize the corresponding *AVR* of *M. oryzae* and triggers the immunity reaction. So far, 12 *AVR* genes in *M. oryzae* have been cloned: *AVR-Pi54*[4], *AVR-Pi9*[5], *AVR-Pib*[6], *AVR-Pia*[7], *AVR-Pii*[7], *AVR-Pik/km/kp*[7], *AVR-Pizt*[8], *ACE1*[9], *AVR-Pita*[10], *AVR1-CO39*[11], *PWL1*[12], and *PWL2*[13]. The *AVR-Pik/km/kp* gene of *M. oryzae* inspects the effectiveness of *R* gene *Pik/km/kp. AVR-Pik/km/kp* encodes a putative secreted protein with 113 amino acids containing two conserved motifs: motif-1, [LI]xAR[SE][DSE]; and motif-2, [RK]CxxCxxxxxxxxxxxxH (similarity to the C2H2 zinc finger motif). [7] The *AVR-Pik/km/kp* gene was cloned from the isolate of Ina168 but absent in the assembled sequence of isolate 70-15, which is recognized by host resistance of Pik protein and triggers the defense response.[7] Five *AVR-Pik* alleles (*AVR-Pik-A, AVR-Pik-B, AVR-Pik-C, AVR-Pik-D, AVR-Pik-E*) were found[7], and the *AVR-Pik-D* (20.5%) and *AVR-Pik-E* (1.4%) was detected among 77 isolates.[14] Four *AVR-Pik* alleles (*AVR-Pik-A, AVR-Pik-C, AVR-Pik-D, AVR-Pik-E*) were found among 39 isolates from world-wide (three isolates from Europe, six isolates from America, seven isolates from Africa and 23 isolates from Asia), and the *AVR-Pik-D* was the highest frequent allele (15 out 39), while the *AVR-Pik-A, AVR-Pik-C, AVR-Pik-E* alleles had similar frequencies (7-9 out of 39).[15] *AVR-Pik/km/kp* has evolved via gene gain/loss manners[7], while substitution mutations were observed in coding regions of the *AVR-Pik/km/kp* in *M. oryzae* populations and 16 SNPs were found in non-signal domain harbored regions in Chinese rice blast isolates.[16]

The *Pik* locus located on the long arm of chromosome 11, and the resistance function has been reported.[17–20] In the *Pik* locus, five rice blast *R* genes (*Pik, Pik-m, Pik-p, Pik-h* and *Pik-s*) involved, among of which four *R* genes (*Pik, Pik-m, Pik-p* and *Pik-h*) have been isolated,[18,21–24] and *Pik* was regard as a younger allele at the locus[22]. *Pik, Pik-m, Pik-p* and *Pik-h* were cloned and those were encode a putative CC-NBS-LRR protein.[18,23,25–26] The CC domain of *Pik-1* physically binds the *AVR-Pik* effector of *M. oryzae* to trigger Pik-specific resistance.[15,23] The rice resistance gene *Pik-s* is still not cloned. The monogenic lines containing 24 rice blast resistance genes including *Pik, Pik-m, Pik-p, Pik-h* and *Pik-s* were developed, which will used to characterize the pathogenicity of rice blast fungus.[27]

*Pikm* and *Pikp* exhibited a high level of resistance to blast fungus from Fujian Province, and can used as resistance breeding parents in Fujian Province[28]. *Pikm, Piks*, and *Pikp* were moderate resistant in Sichuan and Guizhou Provinces, China.[29] *Pikm, Piks*, and *Pik* were moderate resistant, while *Pikh* exhibited high resistance in Guangdong Province, China[30], and 35.4% of 82 rice germplasm resources carrying *Pikh* detected by molecular.[31] While there were 80 carrying *Pik* locus in 229 rice cultivars and breeding material from Fujian Province based on PCR detection.[32] The different resistance spectrum of *Pik, Pikm, Pikp, Pikh* and *Piks* involved in the *Pik* locus were detected by 282 blast isolates collected from Yunnan Province, China.[33] The *R* genes of *Pik* locus exhibit high resistance to Chinese rice blast fungus.

Further understanding the molecular evolution of *AVR* gene has potential implications for the development of resistance breeding and the rational use of resistance genes in production, and the deployment of more effective strategies to control disease. Among long period interaction between the pathogen and its host, the hosts apply the resistance genes to prevent infection by the pathogen, on the other hand, the pathogen attempts to overcome them, and the co-evolution of pathogen and its host was discernible at the genome level.[34–35] The pathogen via mutation to adapt the host novel alleles and environment, while the pathogen genome structure is strongly varied and impacted under host selection.[15,36–37]

The goal of the present study was to analyze DNA sequence variation of *AVR-Pik/km/kp* alleles in field isolates of *M. oryzae* in order to understand the variation and co-evolution mechanism of *M. oryzae AVR-Pik/km/kp* alleles and rice *Pik* alleles in Yunnan Province.

## Results

### The Efficacy of *Pik* loci genes and detection frequency of *AVR-Pik* allele

Based on the disease reactions, the efficacy of *Pik* loci genes of *Pik, Pikm, Pikp, Pikh* and *Piks* were examined. Some 223, 256, 154, 276 and 83 out of the 366 isolates tested were avirulent to the *Pik, Pikm, Pikp, Pikh* and *Piks* gene containing rice monogenic line IRBLk-K, IRBLkm-Ts, IRBLkp-K60, IRBLkh-K3 and IRBLks-F5, respectively (Table 1), the avirulence frequency to *Pik, Pikm, Pikp, Pikh* and *Piks* were 60.9, 69.9, 42.1, 75.4 and 22.7%, respectively; while the remaining 143, 110, 212, 90 and 283 isolates were virulent to the corresponding *R* gene (Table 1). Out of 366 isolates, 278 *AVR-Pik/km/kp* alleles were amplified by *AVR-Pik/km/kp (AVR-Pik* allele) specific primers (pex31F/pex31R) (Table 1), the mean percentage of *AVR-Pik/km/kp* allele was 76.0%. The highest percentage of *AVR-Pik/km/kp* was 90.3% in *M. oryzae* population collected from north-eastern Yunnan, whereas the lowest was 66.7% from north-western Yunnan (Table 1). The percentages of *AVR-Pik/km/kp* were 77.8, 90.3, 66.7, 72.7, 89.3 and 68.3% in central, north-eastern, north-western, south-eastern, south-western and western Yunnan, respectively. Similarly, the percentages of *AVR-Pik/km/kp* were 74.5 and 77.0% in *Xian/Indica* (*XI*) and *Geng/Japonica* (*GJ*) rice growing regions in Yunnan. These findings suggest that *Pik* loci have different effective use in preventing infections by blast in most rice production areas in Yunnan.

**Table 1.**
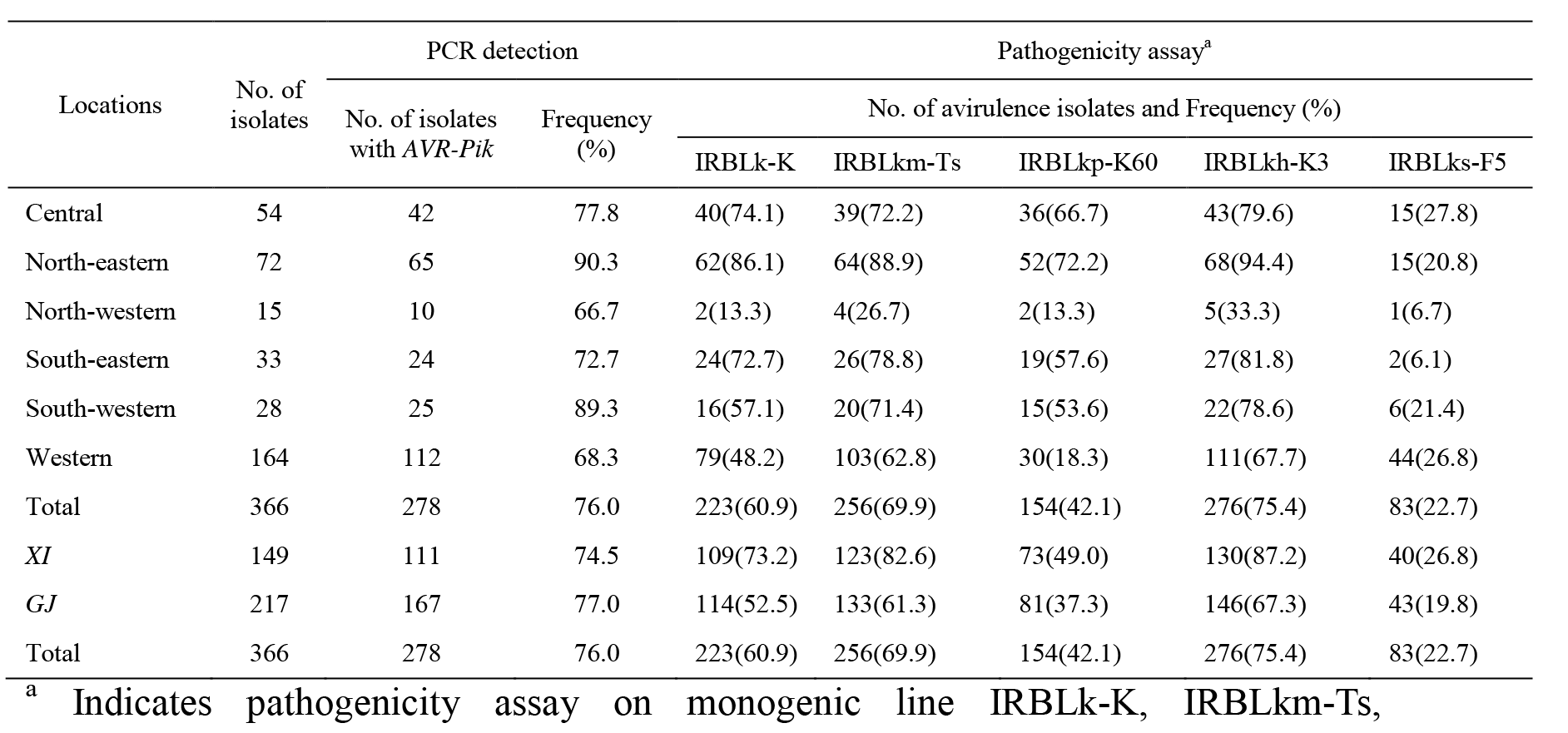
Distribution of *AVR-Pik* genes and avirulence isolates of *M. oryzae* collected from Yunnan, China to IRBLk-K, IRBLkm-Ts, IRBLkp-K60, IRBLkh-K3

| Locations | No. of isolates | PCR detection |  | Pathogenicity assay<sup>a</sup> |  |  |  |  |
| --- | --- | --- | --- | --- | --- | --- | --- | --- |
|  |  | No. of isolates with *AVR-Pik* | Frequency (%) | No. of avirulence isolates and Frequency (%) |  |  |  |  |
|  |  |  |  | IRBLk-K | IRBLkm-Ts | IRBLkp-K60 | IRBLkh-K3 | IRBLks-F5 |
| Central | 54 | 42 | 77.8 | 40(74.1) | 39(72.2) | 36(66.7) | 43(79.6) | 15(27.8) |
| North-eastern | 72 | 65 | 90.3 | 62(86.1) | 64(88.9) | 52(72.2) | 68(94.4) | 15(20.8) |
| North-western | 15 | 10 | 66.7 | 2(13.3) | 4(26.7) | 2(13.3) | 5(33.3) | 1(6.7) |
| South-eastern | 33 | 24 | 72.7 | 24(72.7) | 26(78.8) | 19(57.6) | 27(81.8) | 2(6.1) |
| South-western | 28 | 25 | 89.3 | 16(57.1) | 20(71.4) | 15(53.6) | 22(78.6) | 6(21.4) |
| Western | 164 | 112 | 68.3 | 79(48.2) | 103(62.8) | 30(18.3) | 111(67.7) | 44(26.8) |
| Total | 366 | 278 | 76.0 | 223(60.9) | 256(69.9) | 154(42.1) | 276(75.4) | 83(22.7) |
| *XI* | 149 | 111 | 74.5 | 109(73.2) | 123(82.6) | 73(49.0) | 130(87.2) | 40(26.8) |
| *GJ* | 217 | 167 | 77.0 | 114(52.5) | 133(61.3) | 81(37.3) | 146(67.3) | 43(19.8) |
| Total | 366 | 278 | 76.0 | 223(60.9) | 256(69.9) | 154(42.1) | 276(75.4) | 83(22.7) |

<sup>a</sup> Indicates pathogenicity assay on monogenic line IRBLk-K, IRBLkm-Ts, IRBLkp-K60, IRBLkh-K3 and IRBLks-F5 containing *Pik*, *Pikm*, *Pikp*, *Pikh* and *Piks*, respectively. *XI* and *GJ* indicates *Xian/Indica* and *Geng/Japonica* respectively.

IRBLkp-K60, IRBLkh-K3 and IRBLks-F5 containing *Pik, Pikm, Pikp, Pikh* and *Piks*, respectively. *XI* and *GJ* indicates *Xian/Indica* and *Geng/Japonic* respectively.

### Novel of *AVR-Pikh* gene was identified association with *AVR-Pik/km/kp* alleles

The *AVR-Pik/km/kp* gene is an effector gene with 342 nucleotides encoding a putative secreted protein possessing one signal peptide of 57 first nucleotides in exon in the open reading frame (ORF)[7]. A total of 10 *AVR-Pik* haplotypes, including the five original *AVR-Pik* alleles of *AVR-Pik_D* (GenBank Accession No. AB498875) (H01), *AVR-Pik_A* (AB498876) (H02), *AVR-Pik_B* (AB498877) (H03), *AVR-Pik_C* (AB498878) (H04), and *AVR-Pik_E* (AB498879) (H05) were identified based on the DNA sequence assemblies of 201 isolates (Table 2). The remaining 77 isolates were sequenced, but they had double peaks, and were removed for further analysis. Five novel *AVR-Pik/km/kp* haplotyes of H06-H10 were identified. Alignment of DNA sequence assemblies of the *AVR-Pik/km/kp* gene from 201 isolates revealed that there were six polymorphic sites in the exon region, and none of them in the signal peptid region (Table 2). Six sites in the exon region resulted in amino acid substitutions (Table 3). Moreover, the *AVR-Pik/km/kp* allele sequence assemblies among the 201 isolates were predicted to produce 10 functional proteins (Table 3). Among these 10 proteins, amino acid variations were predicted to occur at five positions. All variations occurred throughout the protein, except of the putative secreted proteins possessing the [RK]CxxCxxxxxxxxxxxxH] motif (Table 3; S1 Fig). Amino acid variations at M78K were found in six isolates, all of which were virulent on monogenic lines IRBLk-K (with *Pik*), IRBLkm-Ts (with *Pikm*), IRBLkp-K60 (with *Pikp*), IRBLkh-K3 (with *Pikh*) and IRBLks-F5 (with *Piks*) (Table 3). This suggests that amino acid 78M is critical for avirulent function of *AVR-Pik/km/kp/kh* loci. The isolates of H01, H07 and H09 haplotypes held the avirulence genes of *AVR-Pik/km/kp/kh*, the isolates of H05 and H08 held *AVR-Pik/km/kh*, the isolates of H06 held *AVR-Pikm/kh*, the isolates of H02 and H03 haplotypes held *AVR-Pikh*, because those isolates were avirulent to corresponding *R* gene(s) (Table 3). While the isolates of H04 and H10 had overcome the resistance of all *Pik* alleles on the loci (Table 3). These findings suggest the novel avirulence gene *AVR-Pikh* was identified, and the evolution of *AVR-Pik* alleles of *M. oryzae* were involved. While the 10 haplotyes did not hold *AVR-Piks*, because of those isolates were virulent to monogenic line IRBLks-F5 (holding *Pi-ks*) (Table 3). Some 75 isolates contained *AVR-Pik/km/kp/kh* (frequency 36.4%), 55 isolates contained *AVR-Pik/km/kh*, (frequency 26.7%), four isolates contained *AVR-Pikm/kh* (frequency 1.9%), 50 isolates contained *AVR-Pikh* (frequency 24.9%). Some 17 isolates did not contain these avirulence gens (S1 Table). In summary, five novel *AVR-Pik* loci were identified, and 91.5% of total isolates contained *AVR-Pikh*, which is widely distributed in south-western China.

**Table 2.**
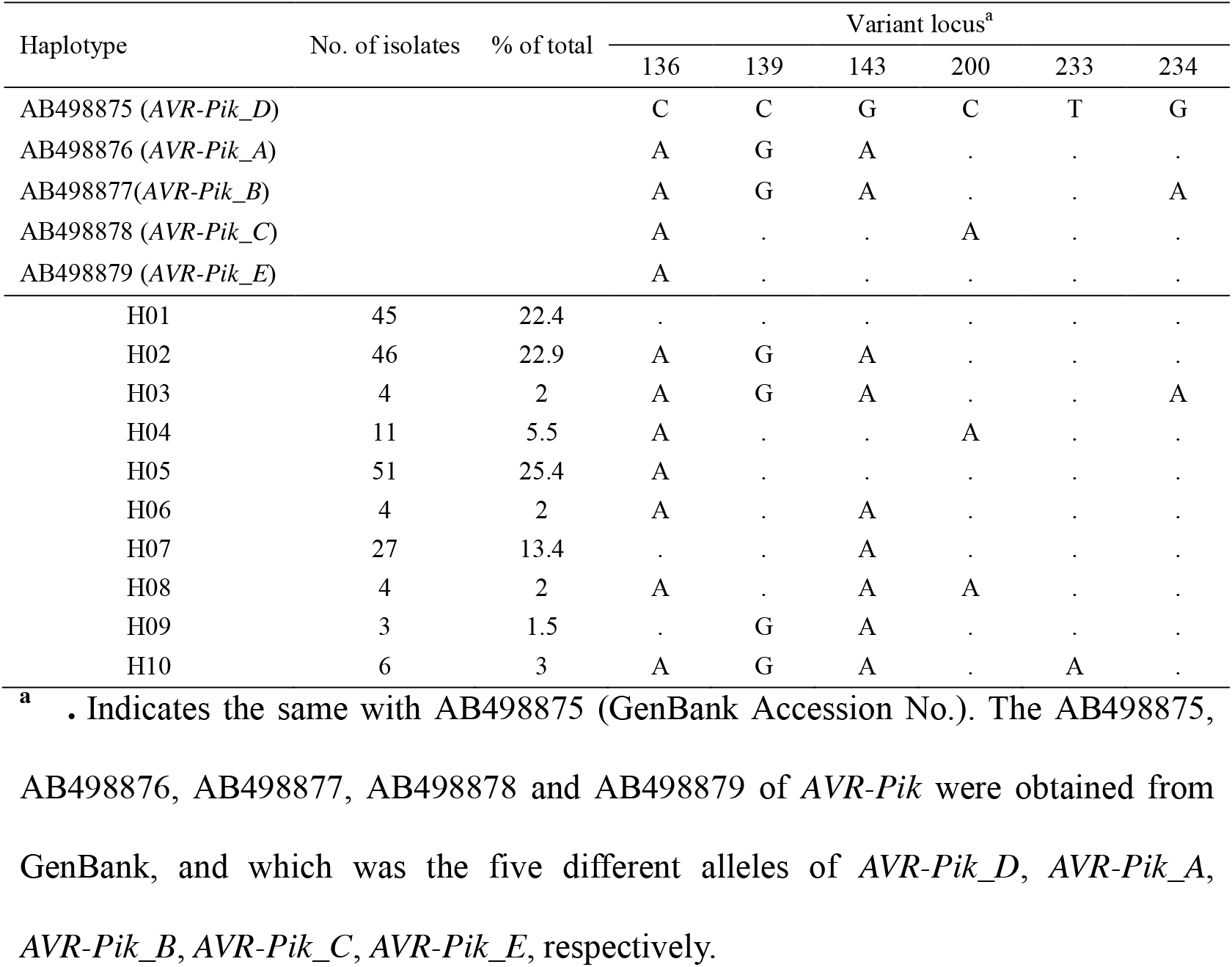
Haplotypes of *AVR-Pik* loci in rice blast fungus of Yunnan, China

| Haplotype | No. of isolates | % of total | Variant locus<sup>a</sup> |  |  |  |  |  |
| --- | --- | --- | --- | --- | --- | --- | --- | --- |
|  |  |  | 136 | 139 | 143 | 200 | 233 | 234 |
| AB498875 (*AVR-Pik_D*) |  |  | C | C | G | C | T | G |
| AB498876 (*AVR-Pik_A*) |  |  | A | G | A | . | . | . |
| AB498877 (*AVR-Pik_B*) |  |  | A | G | A | . | . | A |
| AB498878 (*AVR-Pik_C*) |  |  | A | . | . | A | . | . |
| AB498879 (*AVR-Pik_E*) |  |  | A | . | . | . | . | . |
| H01 | 45 | 22.4 | . | . | . | . | . | . |
| H02 | 46 | 22.9 | A | G | A | . | . | . |
| H03 | 4 | 2 | A | G | A | . | . | A |
| H04 | 11 | 5.5 | A | . | . | A | . | . |
| H05 | 51 | 25.4 | A | . | . | . | . | . |
| H06 | 4 | 2 | A | . | A | . | . | . |
| H07 | 27 | 13.4 | . | . | A | . | . | . |
| H08 | 4 | 2 | A | . | A | A | . | . |
| H09 | 3 | 1.5 | . | G | A | . | . | . |
| H10 | 6 | 3 | A | G | A | . | A | . |

<sup>a</sup> . Indicates the same with AB498875 (GenBank Accession No.). The AB498875, AB498876, AB498877, AB498878 and AB498879 of *AVR-Pik* were obtained from GenBank, and which was the five different alleles of *AVR-Pik\_D*, *AVR-Pik\_A*, *AVR-Pik\_B*, *AVR-Pik\_C*, *AVR-Pik\_E*, respectively.

**Table 3.**
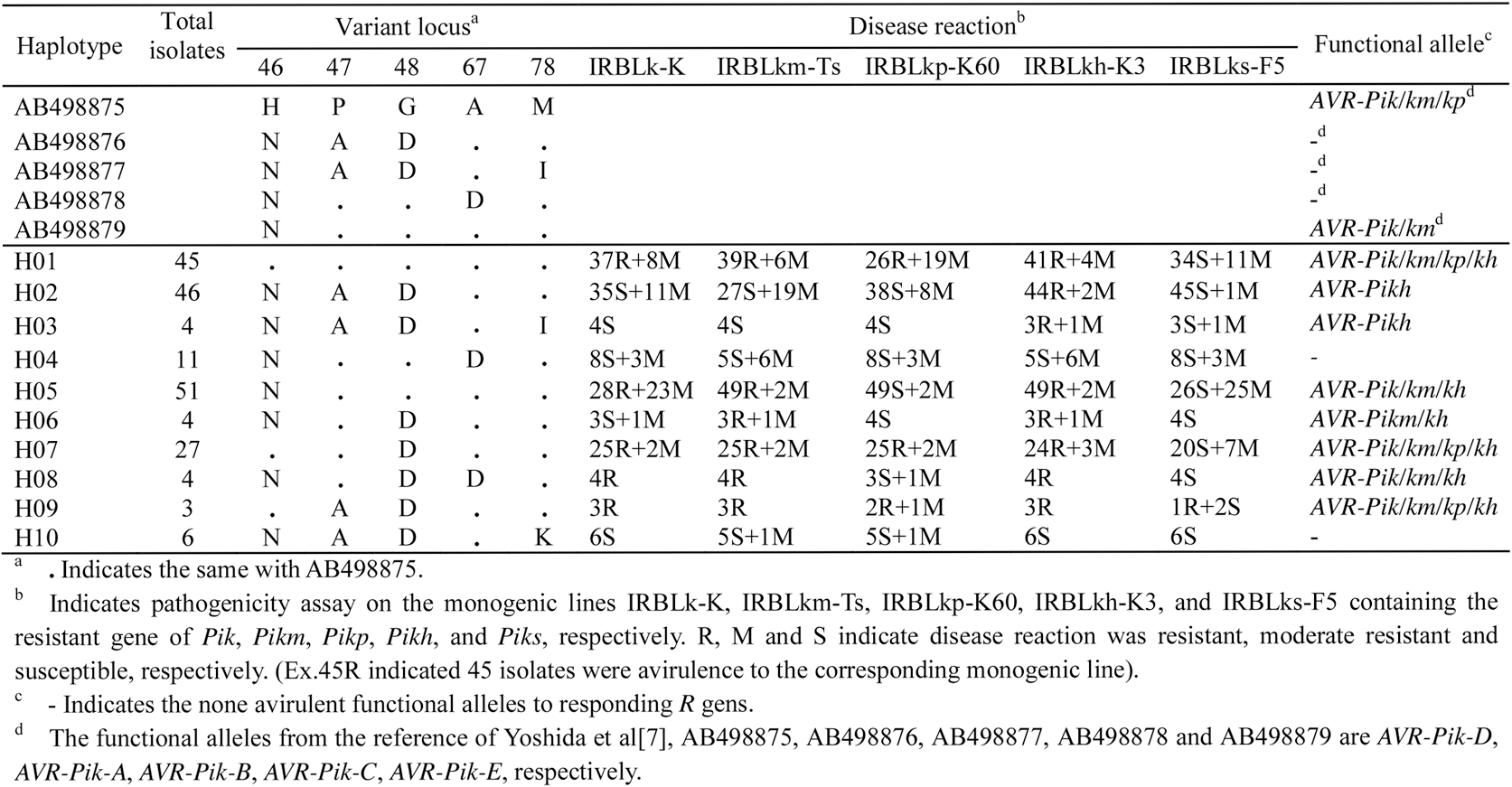
Variation of the *AVR-Pik* loci proteins in rice blast fungus of Yunnan, China

### Stepwise evolution and haplotype diversity of *AVR-Pik* loci in *M. oryzae*

Among 10 *AVR-Pik* haplotypes, the haplotypes of H01, H02, H03, H04 and H05 were identical with the original *AVR-Pik* alleles of *AVR-Pik_D* (GenBank Accession No. AB498875), *AVR-Pik_A* (AB498876), *AVR-Pik_B* (AB498877), *AVR-Pik_C* (AB498878), and *AVR-Pik_E* (AB498879) (Table 2), respectively. Seven haplotypes were detected in 88, 37 and 39 *M. oryzae* from western, central and north-eastern Yunnan, respectively, six haplotypes were detected in 17 *M. oryzae* from south-eastern Yunnan, three haplotypes were detected in 10 *M. oryzae* from south-western Yunnan, and only one haplotype was detected in 10 *M. oryzae* from north-western Yunnan (Table 4). Ten and eight haplotypes were found in the *GJ* and *XI* rice growing regions, and the Diversity Index (DI) was 0.79 and 0.75 for those regions, respectively. Similarly, the DI was 0.78, 0.68, 0.65, 0.62, 0.54, and 0 for north-eastern, central, western, south-eastern, south-western, and north-western Yunnan, respectively (Table 4). In summary, the DI of *AVR-Pik* alleles was ordered in Yunnan Province as: north-eastern>central>western>south-eastern>south-western>north-western. The DI of *AVR-Pik* alleles in *GJ* rice growing region was similar that of *XI*.

**Table 4.**
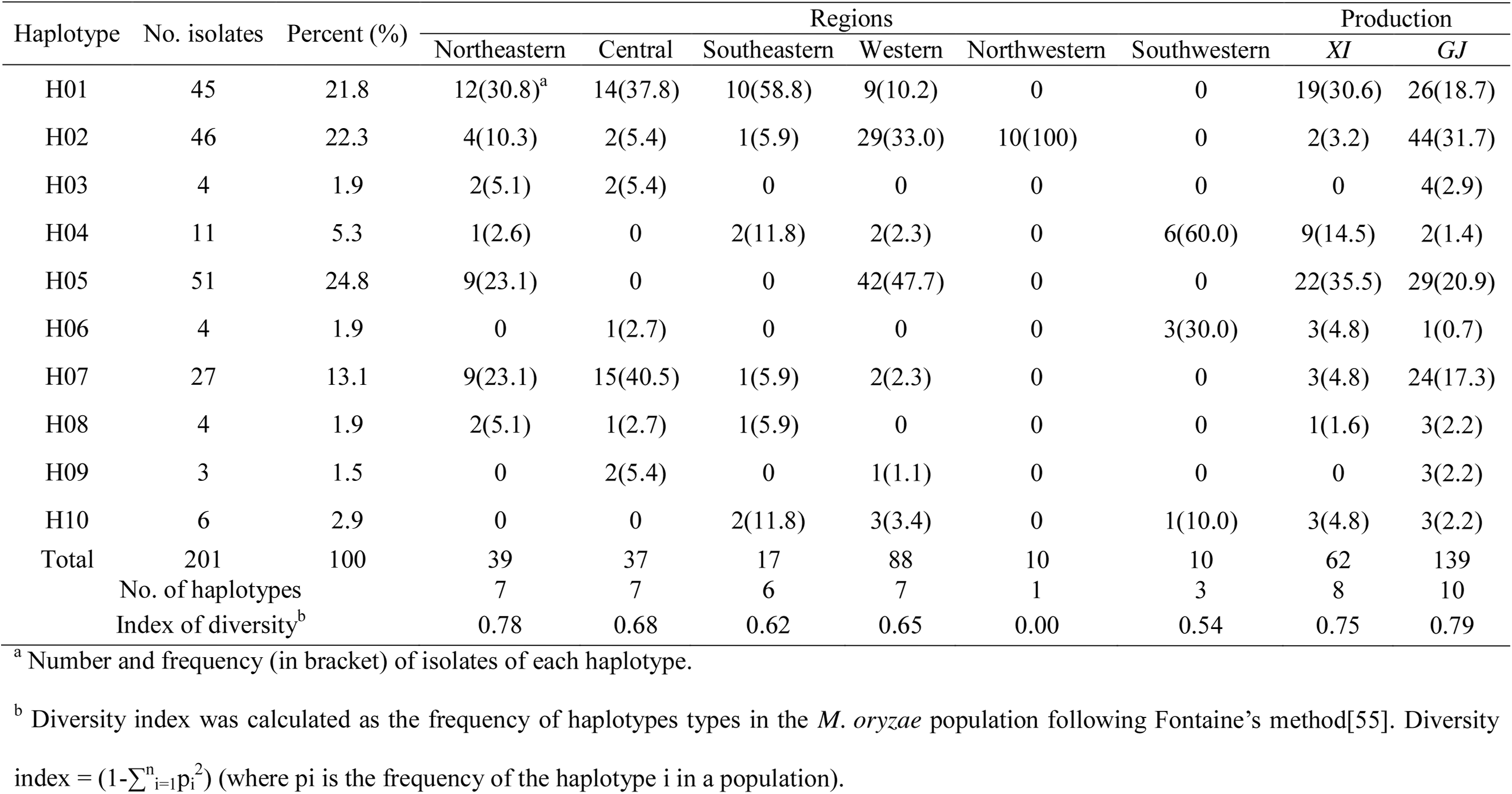
Distribution of *AVR-Pik* haplotype in different rice-growing-regions

Six nucleotide variations in the exons of *AVR-Pik* alleles was observed (S1 Fig and Table 2), and a haplotype network based on sequence variation was developed (Fig 1). Four micro-evolutionary clusters of *AVR-Pik, AVR-Pikm, AVR-Pikp*, and *AVR-Pikh* were observed among 201 field isolates (Fig 1). Five original *AVR-Pik* alleles of *AVR-Pik_D* (H01), *AVR-Pik_A* (H02), *AVR-Pik_B* (H03), *AVR-Pik_C* (H04), and *AVR-Pik_E* (H05) were involved in the networks. The isolates of H01, H05, H07, H08 and H09 were avirulent to IRBLk-K (with Pik), whereas the isolates of H02, H03, H04, H06 and H10 were virulent to *Pik* (Table 3; Fig 1). The isolates of H01, H05, H06, H07, H08 and H09 were avirulent to IRBLkm-Ts (with *Pikm*), whereas the isolates of H02, H03, H04, and H10 were virulent to *Pikm* (Table 3; Fig 1). The isolates of H01, H07 and H09 were avirulent to IRBLkp-K60 (with Pikp), whereas the isolates of H02, H03, H04, H05, H06, H08 and H10 were virulent to *Pikp* (Table 3; Fig 1). The isolates of H01, H02, H03, H05, H06, H07, H08 and H09 were avirulent to IRBLkh-K3 (with Pikh), whereas the isolates of H04 and H10 were virulent to *Pikh* (Table 3; Fig 1). These findings suggest that there were four distinct stepwise evolved patterns of *AVR-Pik, AVR-Pikm, AVR-Pikp*, and *AVR-Pikh* in rice growing regions of Yunnan.

**Fig 1.**
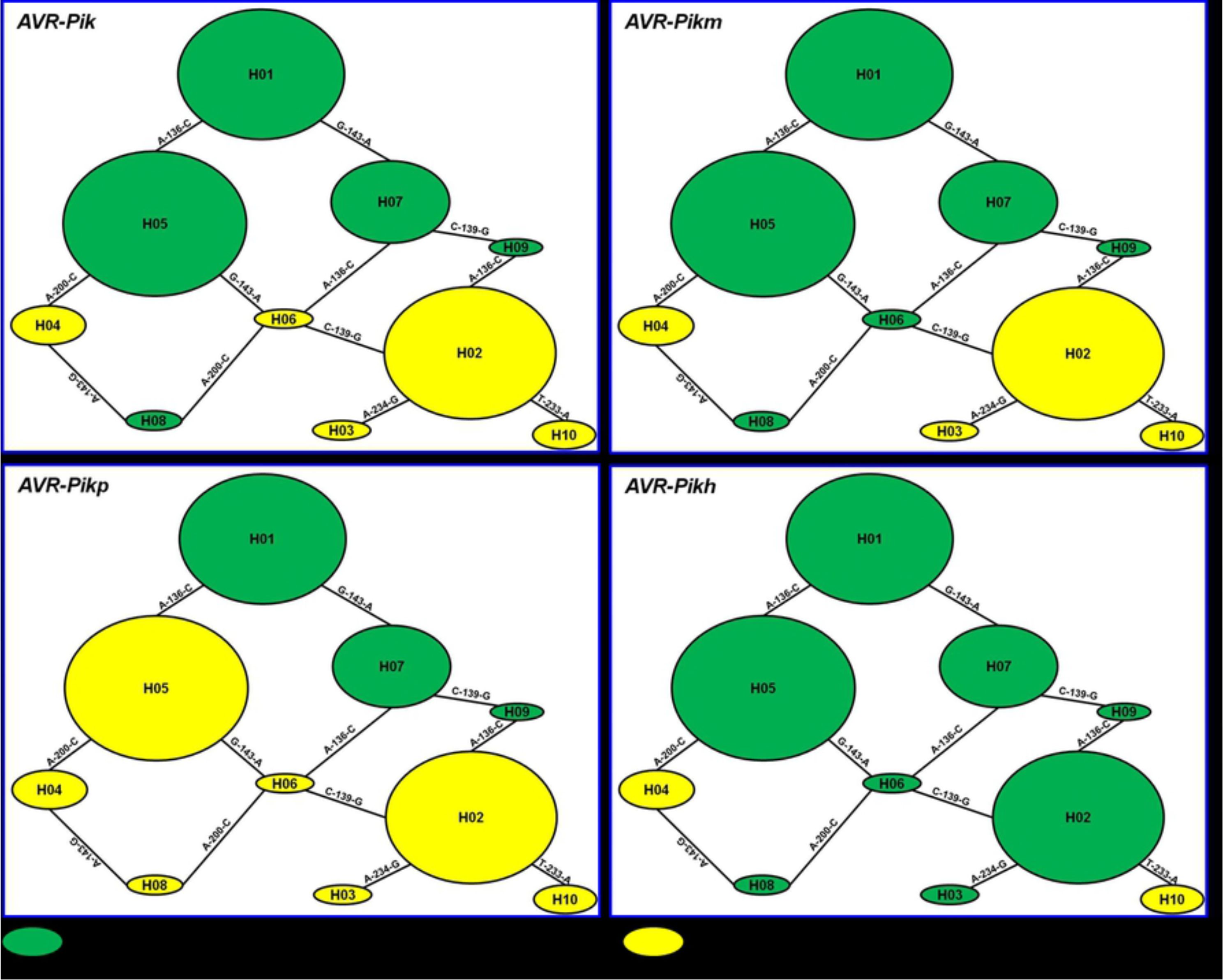
The haplotype network for the 10 *AVR-Pik* alleles. The original *AVR-Pik* allele was designated as the H01 haplotype in the network. Each haplotype was separated by mutational events. All haplotypes were displayed as circles. The size of the circles corresponds to the haplotype frequency. The haplotype H01 to H05 were same with the AB498875, AB498876, AB498877, AB498878 and AB498879 (GenBank Accession No.) of *AVR-Pik* was obtained from GenBank. Green color indicates avirulent to the corresponding *R* gene, yellow color indicates virulent to the corresponding *R* gene.

The possible scenario for *M. oryzae AVR-Pik* alleles-rice *Pik* alleles interactions and co-evolution were constructed (Fig 2). The *AVR-Pik* homolog H01 (*AVR-Pik-D*) were derived from an ancestral *M. oryzae* gene. The *Pik* allele, *Piks*, cannot recognize three alleles *AVR-Pik-D* (H01), H07 and H09, thus the other *Pik* allele, *Pikp*, evolved that can recognize three alleles *AVR-Pik-D* (H01), H07, H09, while the altered alleles H05 (*AVR-Pik-E*) and H08 were evolved to virulence from avirulent origins via nucleotide substitution to avoid recognition by *Pikp* (Table 2; Fig 2). For this situation, another *Pik* allele, *Pik*, evolved that can recognize five alleles *AVR-Pik-D* (H01), H07, H09, *AVR-Pik-E* (H05) and H08. Then, yet another *AVR-Pik* allele, H06, was derived that cannot be recognized by *Pikp* and *Pik*. Next, the rice *R* gene *Pikm* was utilized that recognizes *AVR-Pik-D* (H01), H07, H09, *AVR-Pik-E* (H05), H08 and H06. Then, yet two *AVR-Pik* alleles, *AVR-Pik-A* (H02) and *AVR-Pik-B* (H03), were derived that cannot be recognized by *Pikp, Pik* and *Pikm*. Next, the rice *R* gene *Pikh* was utilized that recognizes *AVR-Pik-D* (H01), H07, H09, *AVR-Pik-E* (H05), H08, H06, *AVR-Pik-A* (H02) and *AVR-Pik-B* (H03). Then another two *AVR-Pik* alleles, *AVR-Pik-C* (H04) and H10, evolved that cannot be recognized by any of the five *Pik* alleles (Table 2; Fig 2). Those showed stepwise evolution of *AVR-Pik* and *Pik* interaction and co-evolution. Interestingly, the *AVR-Pik* allele H07 was derived from H01, that can be recognized by *Pikp, Pik* and *Pikm*. Thus, the altered allele H06 from H07 can avoid recognition by *Pikp* and *Pik*, next the altered allele H08 from H06 can avoid recognition by *Pikp*, while the altered allele H04 from H08 avoids recognition by any of the five *Pik* alleles. Similarly, the H09 derived from H07, that can be recognized by *Pikp, Pik, Pikm* and *Pikh*, thus the altered allele H02 from H09 can avoid recognition by *Pikp, Pik*, and *Pikm* (Table 2; Fig 2). The H05 allele can be recognized by *Pik, Pikm* and *Pikh*, while the altered allele H04 from H05 can avoid recognition by *Pikp, Pik*, and *Pikm* (Table 2; Fig 2). These results suggest that detour evolution of *AVR-Pik* loci of *M. oryzae* were involved during the interaction and co-evolution with the *Pik* loci of *Oryzae* in nature.

**Fig 2.**
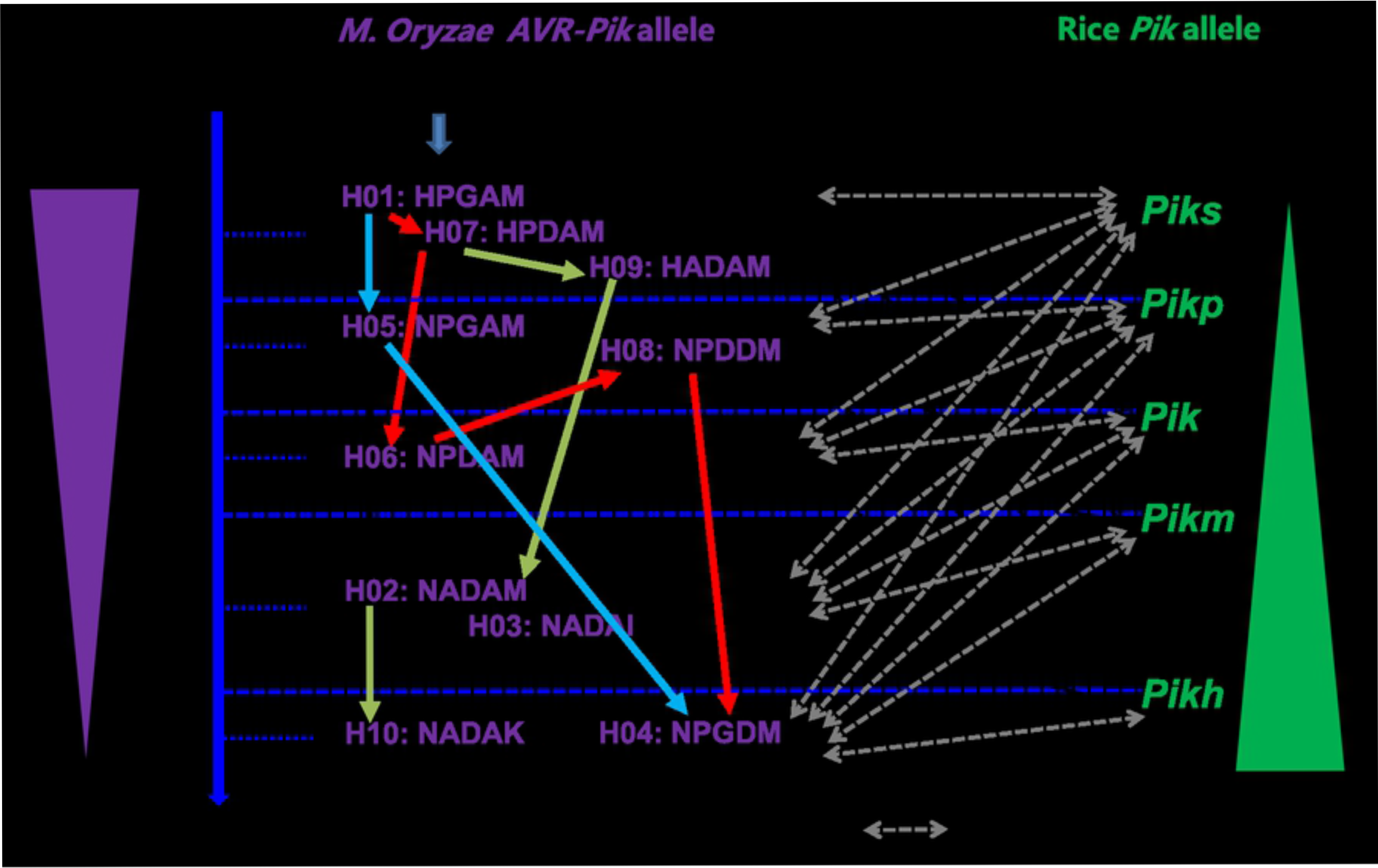
Possible scenario for *M. oryzae AVR-Pik* alleles-rice *Pik* alleles interactions and co-evolution. Chronological order is given on the left (time order). *AVR-Pik* homolog H01 *(AVR-Pik-D*) were derived from an ancestral *M. oryzae* gene. *AVR-Pik-D* (H01), H07 and H09 are recognized by *Pikp*, thus the altered alleles *AVR-Pik-E* (H05) and H08 evolved. In response to this situation, another *Pik* allele, *Pik*, evolved that can recognize five alleles *AVR-Pik-D* (H01), H07, H09, *AVR-Pik-E* (H05) and H08. Then, yet another *AVR-Pik* allele, H06, was derived that cannot be recognized by *Pikp* and *Pik*. Next, the rice *R* gene *Pikm* was utilized that recognizes *AVR-Pik-D* (H01), H07, H09, *AVR-Pik-E* (H05), H08 and H06. Then, yet two *AVR-Pik* alleles, *AVR-Pik-A* (H02) and *AVR-Pik-B* (H03), were derived that cannot be recognized by *Pikp, Pik* and *Pikm*. Next, the rice *R* gene *Pikh* was utilized that recognizes *AVR-Pik-D* (H01), H07, H09, *AVR-Pik-E* (H05), H08, H06, *AVR-Pik-A* (H02) and *AVR-Pik-B* (H03). Then other two *AVR-Pik* alleles, *AVR-Pik-C* (H04) and H10, evolved that cannot be recognized by any of the five *Pik* alleles.

The virulent isolates of H04 and H10 were identified to *Pik* loci (*Pik, Pikm, Pikp, Pikh, Piks*) in most of regions, including north-eastern, south-eastern, south-western, and western Yunnan (Table 3–Table 4; Fig 1). These results suggest that the virulent evolution of *AVR-Pik* loci occurred in most rice-producing regions of Yunnan.

The H01, H04, H05, H06 and H10 haplotypes were mainly distributed in *XI* rice growing regions (Table 4) and H02 and H07 were mainly distributed in *GJ* rice growing regions; whereas H08 was almost equally distributed in *XI* and *GJ* rice growing regions. H03 and H09 were particularly distributed in *GJ* rice growing regions. These findings suggest that the *AVR-Pik* allele in *M. oryzae* was very specifically selected by rice varieties and environments.

### Selection Pressure of *AVR-Pik* in *M. oryzae*

To determine the natural selection pressure of *AVR-Pik* in *M. oryzae* in Yunnan, the Tajima's Neutrality of *AVR-Pik* in *M. oryzae* was tested based on 201 *AVR-Pik* DNA sequences, and the Tajima's *D* was 1.19854 (Table supplement 2). The result suggests that *AVR-Pik* maybe under either strong population expansion or positive selection. The calculation results of three positive-selection models were highly consistent (Fig 3). The sliding window shows the distribution of the Ka/Ks values across all 113 amino acids under the M8, M8a, and M7 models (Fig 3). The results show that the Ka/Ks value of 46^th^, 47^th^, 48^th^, 67^th^ and 78^th^ sites was >1, suggesting that these sites were potentially subjected to purifying selection. The positive selection sites were only observed in the mature protein region among 201 *M. oryzae* isolates with *AVR-Pik* (Fig 3). These results showed that the amino acid sequence was conserved in the signal peptide compared with divergent mature protein region of *AVR-Pik* in *M. oryzae*.

**Fig 3.**
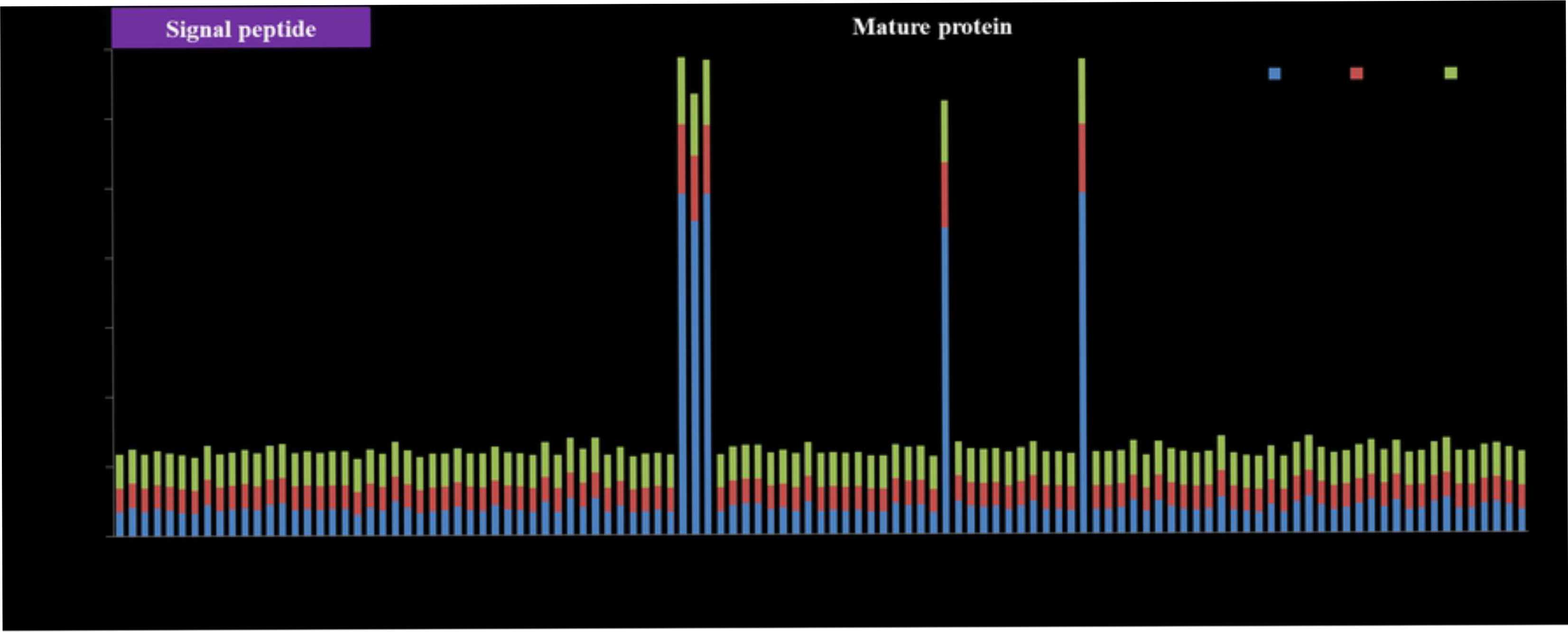
Sliding window of positive-selection sites of the AVR-Pik alleles under M8, M8a, and M7 models. The Y-axis indicates the ratio of the rate of nonsynonymous substitution (Ka) to the rate of synonymous substitution (Ks) (Ka/Ks); the X-axis indicates the position of the AVR-Pik amino acids in the site. The signal region of the variant structure is purple and the black area represents the mature protein region on the label on top of the Figure.

To confirm the resistance performance of alleles of *Pik* in the field, we assayed the seedling and panicle blast disease with monogenic lines carrying the *Pik, Pikm, Pikp, Pikh* which were developed by JIRCAS and IRRI in fields in Mangshi, Lufeng and Yiliang Counties, respectively, in 2015 (Table supplement 3). The result show that IRBLkm-Ts (with *Pikm*), IRBLkp-K60 (with *Pikp*), and IRBLkh-K3 (with *Pikh*) were resistant, while IRBLks-F5 (with *Piks*), IRBLk-Ka (with *Pik*) were susceptible in Mangshi County (Table supplement 3). These results suggest that *M. oryzae* of isolate population holding *AVR-Pikm/kp/kh*. IRBLkh-K3 (with *Pikh*) was resistant in Lufeng and Yiliang, and monogenic lines of IRBLks-F5 (with *Piks*), IRBLk-Ka (with *Pik*), IRBLkm-Ts (with *Pikm*) and IRBLkp-K60 (with *Pikp*) were susceptible in Lufeng and Yiliang Counties, suggesting that *M. oryzae* isolate population holding *AVR-Pikh*. These results are consistent with PCR detection and pathogenicity assays.

## Discussion

In this study, we found five new haplotypes in the *AVR-Pik* DNA sequences among field isolates of *M. oryzae* from various rice-producing regions in Yunnan. Numerous virulence isolates to the *Pik* gene containing rice varieties were identified in field isolates collected in Yunnan, suggesting that *Pik* was defeated in some rice production areas due to extensive development of *Pik* in China. The *Pik* alleles have been deployed and display high rice blast resistance in China.[20,22,32,38] Complete deletions have occurred in *AVR-Pik* sequences among field isolates of *M. oryzae* from various rice-producing countries[15–16], which accords with our results. Numerous isolates inspected from commercial rice fields, containing *AVR-Pik* suggest that *Pik* has effective in preventing rice blast disease. In Yunnan, rice cultivars with *Pikh, Pikp, Pikm, Piks, Pik* were resistant to 81.7, 62.8, 51.9, 43.4 and 39.4% of isolates (282 isolates), respectively[33]. Corresponding values from 146 isolates from Guandong Province were 88.4, 39.0, 0, 1.4 and 57.5%, respectively. [39] These results suggest that some *Pik* alleles has limited effects in these rice production areas. Continued analysis of *AVR-Pik* alleles in these isolates will help us understand the evolutionary mechanism of *AVR-Pik*, and to predict the stability and effectiveness of *Pik* alleles mediated resistance under natural conditions.

Effective variations of DNA sequence were observed in several *AVR* genes (*AVR-Pita1, AVR-Pia*, and *AVR-Pii*) in the telomere regions.[7,40–41] The transposable element insertion in the last exon of the *ACE1* gene[9] and Pot3 inserted in *AVR-Pizt* and *AVR-Pita1* all resulted in new virulent alleles. Based on the DNA sequence analysis,[8,42–43] four variations of point mutation, segmental deletion, complete absence (6.7%) and transposable element (TE) insertion were found in *AVR-Pib*, all of which result in losses of the avirulence function.[6] Three distinct expression profiles were found among seven functional of 16 nucleotide polymorphisms in the *AVR-Pib* genes.[6] These findings showed that *M. oryzae* uses the transposons to change the expression of *AVR* genes in defeating *R* genes. In the present study, the *AVR-Pik* gene was present in most blast populations (76.0%) in Yunnan (Table 1), which was similar to rice blast isolates in Hunan Province.[44] We found significantly more nucleotide variation in the protein coding region of *AVR-Pik* alleles, resulting in changes of amino acids suggesting that there exists intense selection pressure on *AVR-Pik* alleles in Yunnan.

DNA sequence variation was found in exon regions of *AVR-Pik*, and a total of 10 haplotypes were identified based on the six variant nucleotides among 201 isolates collected from Yunnan (Table 2). Five novel variant amino acids of the *AVR-Pik* loci variants in the 201 isolates were identified in the present study, leading to finding five new haplotypes. Based on the virulence analysis of the strains harboring this variation, haplotypes H01, H02, H05 and H07 are more frequent in the field isolates. This probably suggests that loss of these haplotypes may have a larger fitness penalty than others alleles in the *M. oryzae* population. These new alleles allow us to construct a more holonomic network among different alleles of *AVR-Pik* and some novel haplotypes were found. We also identified the putative secreted proteins possessing the [LI]xAR[SE][DSE] and [RK]CxxCxxxxxxxxxxxxH] motif in 201 isolates with *AVR-Pik* alleles (Table 3) which was consistent with the results of Yoshida et al[7]. Some 126, 59, 94 and 15 isolates are variations at the amino acid position of H46N, P47A, G48D, A67D, respectively, and four and six isolates are variations in the amino acid position of M78I and M78K, respectively (Table 3). These results showed the amino acid position of 46^th^, 47^th^, 48^th^, 67^th^ and 78^th^ were the hot variation amino acid sites among proteins of AVR-Pik/km/kp/kh.

During the long co-evolution of plants and pathogens, the pathogen *AVR* genes are recognized by the cognate plant *R* genes in trigging effective defense responses. The divergences of *AVR* genes of the pathogen were shaped by host *R* genes and changing environmental conditions. We observed that the Diversity Index of *AVR-Pik* in *XI* and *GJ* regions were similar (Table 4) and variations of *AVR-Pik* were different between *XI* and *GJ* growing regions (Table 4). These results suggest that adaptive variations have occurred in commercial rice fields in Yunnan.

Yunnan is one of the diversification centers of the cultivated Asian rice *Oryza sativa*. Three wild species of *O. refipogon, O. officinalis* and *O. meyeriana* existed in the area.[45] Over 5000 accessions of rice germplasms were collected from fields and preserved. Among them, 227 rice accessions were characterized by a set of differential rice blast isolates, and 38 and 25 of 227 rice accessions contained the rice blast resistance genes *Pik* and *Pikm*, respectively.[45] While the neutrality test of Tajima's *D* was 1.19854 (Table supplement 2), it suggests *AVR-Pik/km/kp/kh* loci may be under population expansion or purifying selection shaped by the cognate gene *Pik* loci in rice growing regions of Yunnan. Most of isolates carried *AVR-Pikh* and *Pikh* with high resistance in Yunnan and Guangdong Provinces. This may be due to *Pikh* being a widely distributed resistant gene in rice accessions. These results accord with Zhai et al[22].

The *AVR-Pik* was recognition specificity by *Pik* of rice, and AVR-Pik directly specific physically binds the N-terminal coiled-coil domain of Pik. These observations were confirmed by yeast two-hybrid and co-immunoprecipitation in plant assays.[15] Four alleles of *AVR-Pik* (*AVR-Pik_D, AVR-Pik_E, AVR-Pik_A, AVR-Pik_C*) in Japanese isolate populations were in the manner of co-evolution with the rice *Pik* alleles *Pikp, Pik* and Pikm.[15] Four alleles of *AVR-Pik* in Chinese *M. oryzae* population stepwise evolution between rice *Pik* allele *Pikp, Pik, Pikm*, and *Pkh* were found.[16] Highly variable *Pik* alleles were observed, and both stepwise *AVR-Pik* of *M. oryzae* and *Pik* of rice occurred in field conditions.[16] These observations indicate that *AVR-Pik* has been strongly targeted by hosts.[16] In this present study, we found both the detour and stepwise evolved *AVR-Pik* alleles-rice *Pik* alleles interactions and co-evolution (Table 3; Fig 2), which implies high diversity of rice varieties in Yunnan. The *AVR-Pik* alleles have been regularly under selection stress by antagonistic alleles in host populations. Similarly, the wheat-infecting lineages from Brazil and Bangladesh appeared genetically distinct and displayed reticulate evolution on population genomic analyses of transcriptomic single nucleotide polymorphisms. [46]

The stepwise mutation process has been demonstrated for virulence acquisition in *Fusarium oxysporum* f. sp. *Ciceris* and *Puccinia striiformis* f. sp. *Tritici*. [47–49] In the present study, we found one major mutation evolution of *AVR-Pik* allele and seven minor mutation evolution patterns (Fig 2). The alternative mutation pattern can seemingly convert from avirulence to virulence via seldom mutation, and showed higher efficiency (Fig 2). These may be due to the strong positive selection pressure by the corresponding *Pik* allele on the host and environment. Similarly, *AVRL567* can convert from avirulent to virulent by a set of stepwise mutants in amino acid substitution.[50] Stepwise evolved procession has observed in *AVR-Pik*.[15–16] The possible evolution of *AVR-Pik* found in the present study, showed more complex evolution than expected in the rice growing regions of Yunnan.

## Conclusion

We detected five novel haplotypes in the field population by using 201 isolates, constructed a complex network of *AVR-Pik* alleles, and evaluated the effectiveness of *Pik* alleles in rice production areas of Yunnan. Our findings support the premise that functional *AVR-Pik* possesses diversified sequence structures and can avoid recognition by host via multiple site variations. Haplotype H10 originates from frequently distributed H2, H4 jump from H5 and/or H8, can overcome all detected *Pik* alleles to date. Although H4 and H10 haplotypes have low frequency, but surveillance of these two alleles in field populations is crucial because of its high risk of leading to the breakout on the background of *Pik* rice variety. Management must retard the selection stress on the allele, possibly by avoiding its proliferation in agricultural practices. Prediction of blast occurring should be based on the frequency and distribution of allele of multiple loci v.g. *Pik* and *AVR-Pik* in isolate populations in field conditions.

## Materials and methods

### Rice cultivars, fungal isolates, culture, and pathogenicity assays

The *Pik, Pikm, Pikp, Pikh, Piks* gene containing rice monogenic line IRBLk-K, IRBLkm-Ts, IRBLkp-K60, IRBLkh-K3 and IRBLks-F5, respectively, and the susceptible backcrossing parent Lijiangxintuanheigu (LTH without *Pik*) were used for pathogenicity assays. A total of 366 isolates were collected, single spore purified, and examined. All isolates were stored at −20°C on filter paper and grown at room temperature under blue and white fluorescent lighting on petri dishes containing oatmeal agar for spore production. Disease reactions were determined using a modified standard pathogenicity assay, as described by Jia et al.[51] Specifically, rice seedlings at the 3- to 4- leaf stage were placed in a plastic bag and were spray inoculated with a spore suspension of 1-5×10^5^ spores/mL. After inoculation, the plastic bags were sealed to maintain high relative humidity (90-100%) for 24 h before removing the plants from bags. Subsequently, plants were maintained in a greenhouse for an additional six days, to allow the development of disease symptoms. The disease reactions were rated based on visual number and amount of lesions at the second youngest leaf using the 0-5 disease scale. A value of 0-1 is resistant, 2 is moderately resistant and 3-5 is susceptible. Five seedlings were used each time and the experiment was repeated one more time, and the mean disease scores were used to determine resistance versus susceptibility.

### DNA preparation, PCR amplification, and DNA sequencing

Fungal isolates were grown in complete liquid media at 25°C for six-eight days to produce mycelium under dark conditions. DNA was then isolated from mycelia using the CTAB method.[52] Primers pex31F (5’-TCGCCTTCCCATTTTTA-3’) and pex31R (5’-GCCCATGCATTATCTTAT-3’) were used to amplify the *AVR-Pik* allele and for sequencing using the methods of Yoshida et al.[7] Specifically, PCR reactions were performed using 2×Taq PCR MasterMix (Tiangen Biotech Co. Ltd., Beijing, China). Each PCR reaction consisted of the following components: 25 μl of Taq PCR Master Mix (contains 25U of Taq DNA polymerase, 10X Tiangen PCR buffer, 15 mM MgCl2, and 200 μM of each dNTP), 1 μl of each 10 μM primer, 2 μl of fungal genomic DNA, and 21 μl distilled water (provided by the Tiangen Kit). Reactions were performed in a BIO-RAD Thermal Cycler (C1000, Bio-Rad Laboratories, Life Science Research, Hercules, CA, USA) with the following PCR program: 1 cycle at 95°C for 3 min for initial denaturation, followed by 29 cycles at 95°C for 30 s, 60°C for 30 s, 72°C for 30 s, and a final denaturation of 72°C for 7 min. All PCR reactions were repeated three times (20 μl for detection, 50 μl for sequencing). The size of the amplified fragment was estimated by DL2000 DNA Ladder (Tiangen Biotech Co. Ltd., Beijing, China). PCR products were sequenced using the same primers as mentioned above for PCR amplification. DNA was sequenced by Shanghai Life Technologies Biotechnology Co., Ltd. (Shanghai, China). The amplicon from each isolate was sequenced three times.

### Resistance evaluation of *Pik* alleles in field

The monogenic lines of IRBLk-Ka, IRBLkm-Ts, IRBLks-F5, IRBLkp-K60, IRBLkh-K3 (carrying the *Pik, Pikm, Piks, Pikp, Pikh*, respectively.), which were planting in fields in Mangshi, Lufeng and Yiliang Counties of Yunnan Province, respectively, in 2015. The seedling and panicle blast disease were surveyed, and the resistance were evaluated.

### Data analysis

DNA sequences of *AVR-Pik* were assembled by Vector NTI software Suite V.10 (Invitrogen, Carlsbad, California, USA) and aligned using DNASTAR V7.10 software (http://www.dnastar.com/). The number of DNA haplotypes, polymorphic sites (*π*), and the sliding window were calculated using DnaSP v5.10.01 software.[53] Haplotype network analysis was performed using TCS1.21 (http://darwin.uvigo.es/). [54]Diversity Index was calculated as the frequency of haplotypes or protein types in the rice blast fungus population following the method of Fontaine et al.[55] Diversity 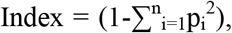 where pi is the frequency of the haplotype i in a population. Tajima's neutrality test was performed using MEGA V5.10.

### Supporting information

**S1 Fig. Diversification of *AVR-Pik* in avirulent isolates**. Distribution of variation of the *AVR-Pik* alleles was analyzed using sliding window. X-axis shows the distribution of variation within the full region, including signal peptide and exon of *AVR-Pik*. Lower pane indicates the corresponding schematic presentation of the signal peptide and exon of *AVR-Pik*. Window length: 1; Step size: 1. π value corresponds with the level of variation at each site because it is the sum of pair-wise differences divided by the number of pairs within the population.

**S1 Table**. Distribution of *AVR-Pik* loci in rice blast fungus.

**S2 Table**. Tajima's Neutrality Test of *AVR-Pik* in M. *oryzae*. The analysis involved 201 nucleotide sequences of *AVR-Pik. m* indicates number of sequences, *S* indicates number of segregating sites, *Ps* indicates *S/n, Θ* indicates p_s_/a_1_, *π* indicates nucleotide diversity, and *D* is the Tajima test statistic. Tajima's D: 1.19854, Statistical significance: Not significant, P>0.10.

**S3 Table**. Summary of disease reaction of monogenic lines with *Pik* alleles in fields. The pathogenicity assay on the monogenic lines IRBLk-K, IRBLkm-Ts, IRBLkp-K60, IRBLkh-K3, and IRBLks-F5 containing the resistant genes of *Pik, Pikm, Pikp, Pikh*, and *Piks*, respectively. R and S indicate disease reaction was resistant and susceptible, respectively.

## Acknowledgments

The authors gratefully thank Professor Michael A Fullen (The University of Wolverhampton) for useful discussions and proofreading this manuscript. This work was supported by the National Natural Science Foundation of China (31460454) and the Department of Sciences and Technology of Yunnan Province, China (2015HB076 and 2017FA013) to Jinbin Li, and the National Key R&D Program of China (2017YFD0200400).

## References

1. Woolhouse M, Webster J, Domingo E, Charlesworth B, Levin B. Biological and biomedical implications of the co-evolution of pathogens and their hosts. Nature Genetics. 2002; 32(4):569–577.

2. Ma J, Lei C, Xu X, Hao K, Wang J, Cheng Z, et al. *Pi64*, encoding a novel CC-NBS-LRR protein, confers resistance to leaf and neck blast in rice. Molecular Plant-Microbe Interactions. 2015; 28(5):558–568.

3. Deng Y, Zhai K, Xie Z, Yang D, Zhu X, Liu J, et al. Epigenetic regulation of antagonistic receptors confers rice blast resistance with yield balance. Science. 2017; 355(6328):962–965.

4. Ray S, Singh PK, Gupta DK, Mahato AK, Sarkar C, Rathour R, et al. Analysis of *Magnaporthe oryzae* genome reveals a fungal effector, which is able to induce resistance response in transgenic rice line containing resistance gene, *Pi54*. Frontiers in Plant Science. 2016; 7:1–16. doi: 10.3389/fpls.2016.01140

5. Wu J, Kou Y, Bao J, Li Y, Tang M, Zhu X, et al. Comparative genomics identifies the *Magnaporthe oryzae* avirulence effector *AvrPi9* that triggers *Pi9*-mediated blast resistance in rice. New Phytologist. 2015; 206:1463–1475.

6. Zhang S, Wang L, Wu W, He L, Yang X, Pan Q. Function and evolution of *Magnaporthe oryzae* avirulence gene *AvrPib* responding to the rice blast resistance gene *Pib*. Scientific Reports. 2015; 5:11642. doi: 10.1038/srep11642.

7. Yoshida K, Saitoh H, Fujisawa S, Kanzaki H, Matsumura H, Yoshida K, et al. Association genetics reveals three novel avirulence genes from the rice blast fungal pathogen *Magnaporthe oryzae*. Plant Cell, 2009; 21(5):1573–1591.

8. Li W, Wang B, Wu J, Lu G, Hu Y, Zhang X, et al. The *Magnaporthe oryza* avirulence gene *AVR-Pizt* encodes a predicted secreted protein that triggers the immunity in rice mediated by the blast resistance gene *Piz-t*. Molecular Plant-Microbe Interactions. 2009; 22(4):411–420.

9. Fudal I, Bohnert HU, Tharreau D, Lebrun MH. Transposition of MINE, a composite retrotransposon, in the avirulence gene *ACE1* of the rice blast fungus *Magnaporthe grisea*. Fungal Genetics and Biology. 2005; 42(9):761–772.

10. Orbach MJ, Farrall L, Sweigard JA, Chumley FG, Valent B. A telomeric avirulence gene determines efficacy for the rice blast resistance gene *Pi-ta*. Plant Cell. 2000; 12(11):2019–2032.

11. Farman ML, Leong SA. Chromosome walking to the *AVR1-CO39* avirulence gene of *Magnaporthe grisea*: discrepancy between the physical and genetic maps. Genetics. 1998; 150:1049–1058.

12. Kang S, Sweigard JA, Valent B. The *PWL* host specificity gene family in the blast fungus *Magnaporthe grisea*. Molecular Plant-Microbe Interactions. 1995; 8(6):939–948.

13. Sweigard JA. Identification, cloning, and characterization of *PWL2*, a gene for host species specificity in the rice blast fungus. Plant Cell. 1995; 7:1221–1233.

14. Selisana SM, Yanoria MJ, Quime B, Chaipanya C, Lu G, Opulencia R, et al. Avirulence (*AVR*) gene-based diagnosis complements existing pathogen surveillance tools for effective deployment of resistance (*R*) genes against rice blast disease. Phytopathology. 2017; 107(6):711–720.

15. Kanzaki H, Yoshida K, Saitoh H, Fujisaki K, Hirabuchi A, Alaux L, et al. Arms race co-evolution of *Magnaporthe oryzae AVR-Pik* and rice *Pik* genes driven by their phyical interactions. Plant Journal. 2012; 72(6):894–907.

16. Wu W, Wang L, Zhang S, Li Z, Zhang Y, Lin F, et al. Stepwise arms race between *AvrPik* and *Pik* alleles in the rice blast pathosystem. Molecular Plant-Microbe Interactions. 2014; 27(8):759–769.

17. Kiyosawa S. With genetic view on the mechanism of resistance and virulence. Japanese Journal of Genetics. 1987; 41:89–92. In Japanese.

18. Ashikawa I, Hayashi N, Yamane H, Kanamori H, Wu J, et al. Two adjacent nucleotide-binding site-leucine-rich repeat class genes are required to confer *Pikm*-specific rice blast resistance. Genetics. 2008; 180(4):2267–2276.

19. Xu X, Hayashi N, Wang C, Kato H, Fujimura T, Kawasaki S. Efficient authentic fine mapping of the rice blast resistance gene *Pik-h* in the *Pik* cluster, using new Pik-h-differentiating isolates. Molecular Breeding. 2008; 22(2):289–299.

20. Wang L, Xu X, Lin F, Pan Q. Characterization of rice blast resistance genes in the *Pik* cluster and fine mapping of the *Pik-p* locus. Phytopathology. 2009; 99(8):900–905.

21. Ashikawa I, Hayashi N, Abe F, Wu J, Matsumoto T. Characterization of the rice blast resistance gene *Pik* cloned from Kanto51. Molecular Breeding. 2012; 30(1):485–494.

22. Zhai C, Lin F, Dong Z, He X, Yuan B, Zeng X, et al. The isolation and characterization of *Pik*, a rice blast resistance gene which emerged after rice domestication. New Phytologist. 2011; 189(1):321–334.

23. Zhai C, Zhang Y, Yao N, Lin F, Liu Z, Dong Z, et al. Function and interaction of the coupled genes responsible for *Pik-h* encoded rice blast resistance. PLoS ONE. 2014; 9(6):e98067. doi:10.1371/journal.pone.0098067

24. Sharma TR, Madhav MS, Singh BK, Shanker P, Jana TK, Dalal V, et al. High-resolution mapping, cloning and molecular characterization of the *Pi-k^h^* gene of rice, which confers resistance to *Magnaporthe grisea*. Molecular Genetics and Genomics. 2005; 274(6):569–578.

25. Chen J, Peng P, Tian J, He Y, Zhang L, Liu Z, et al. *Pik*, a rice blast resistance allele consisting of two adjacent NBS–LRR genes, was identified as a novel allele at the *Pik* locus. Molecular Breeding. 2015; 35:117. https://doi.org/10.1007/s11032-015-0305-6

26. Yuan B, Zhai C, Wang W, Zeng X, Xu X, Hu H, et al. The *Pik-p* resistance to *Magnaporthe oryzae* in rice is mediated by a pair of closely linked CC-NBS-LRR genes. Theoretical and Applied Genetics. 2011; 122(5):1017–1028. doi: 10.1007/s00122-010-1506

27. Tsunematsu H, Yanoria MJT, Ebron LA, Hayashi N, Ando I, Kato H, et al. Development of monogenic lines of rice for blast resistance. Breeding Science. 2000; 50(3):229–234.

28. Du Y, Ruan H, Shi N, Gan L, Yang X, Chen F. Pathogenicity analysis of *Magnaporthe grisea* against major *Pi*-genes and main rice varieties in Fujian Province. Journal of Plant Protection. 2016; 43(3):442–451. In Chinese.

29. Zhang S, Zhong X, Qiao G, Shen L, Zhou T, Peng Y. Difference in virulence of *Magnaporthe oryzae* from Sichuan, Chongqing and Guizhou. Southwest China Journal of Agricultural Sciences. 2017; 30(2):359–365. In Chinese.

30. Yang J, Chen S, Zeng L, Li Y, Chen Z, Zhu X. Evaluation on resistance of major rice blast resistance genes to *Magnaporthe grisea* isolates collected from indica rice in Guangdong Province, China. Chinese Journal of Rice Science. 2008; 22(2):190–196. In Chinese.

31. Xie Q, Guo J, Yang S, Chen Z, Cheng B, Huang Y, et al. Evaluation of blast resistance spectrum and identification of resistance genes in 82 rice germplasm resources. Guangdong Agricultural Science. 2015; 14:31–35. In Chinese.

32. Chen Z, Tian D, Liang T, Chen Z, Hu C, Wang F, et al. Characterization of the genotypes at the rice blast resistance *Pik* locus in 229 rice cultivars and important breeding materials. Fujian Journal of Agriculture Science. 2016; 31(6):553–559. In Chinese.

33. Li J, Li C, Chen Y, Lei C, Ling Z. Evaluation of twenty-two blast resistance genes in Yunnan using monogenetic rice lines. Acta Phytophylacica Sinica. 2005; 32(2):113–119. In Chinese.

34. Raffaele S, Farrer R, Cano L, Studholme D, Maclean D, Thines M, et al. Genome evolution following host jumps in the Irish potato famine pathogen lineage. Science. 2010; 330(6010):1540–1543.

35. Terauchi R, Yoshida K. Towards population genomics of effector-effector target interactions. New Phytologist. 2010; 187(4):929–939.

36. Stukenbrock E, McDonald B. Population genetics of fungal and Oomycete effectors involved in gene-for-gene interactions. Molecular Plant-Microbe Interactions. 2009; 22(4):371–380.

37. Daugherty M, Malik H. Rules of engagement: Molecular insights from host-virus arms races. Annual Review of Genetics. 2012; 46:677–700. doi: 10.1146/annurev-genet-110711-155522

38. Hua L, Wu J, Chen C, Wu W, He X, Lin F, et al. The isolation of *Pi1*, an allele at the *Pik* locus which confers broad spectrum resistance to rice blast. Theoretical and Applied Genetics. 2012; 125(5):1047–1055.

39. Zhu X, Yang Q, Yang J, Lei C, Wang J, Ling Z. Differentiation ability of monogenic lines to *Magnaporthe grisea* in indica rice. Acta Phytopathologica Sinica. 2004; 34(4):361–368. In Chinese.

40. Chuma I, Isobe C, Hotta Y, Ibaragi L, Futamata N, Kusaba M, et al. Multiple translocation of the *AVR-Pita* effector gene among chromosomes of the rice blast fungus *Magnaporthe oryzae* and related species. PLoS Pathog. 2011; 7(7):e1002147. doi: 10.1371/journal.ppat.1002147.

41. Dai Y, Jia Y, Correll J, Wang X, Wang Y. Diversification evolution of the avirulence gene *AVR-Pita1* in field isolates of *Magnaporthe oryzae*. Fungal Genetics and Biology. 2010; 47(12):973–980.

42. Kang S, Lebrun MH, Farrall L, Valent B. Gain of virulence caused by insertion of a Pot3 transposon in a *Magnaporthe grisea* avirulence gene. Molecular Plant-Microbe Interactions. 2001; 14(5):671–674.

43. Zhou E, Jia Y, Singh P, Correll J, Lee FN. Instability of the *Magnaporthe oryzae* avirulence gene *AVR-Pita* alters virulence. Fungal Genetics and Biology, 2007; 44(10):1024–1034.

44. Xing J, Jia Y, Peng Z, Shi Y, He Q, Shu F, et al. Characterization of molecular identity and pathogenicity of rice blast fungus in Hunan Province of China. Plant Disease. 2017; 101(4):557–561.

45. Li J. Breeding of Yunnan rice. In: Jiang Z, editors. Yunnan Rice. Kunming: Yunnan Science and Technology Press; 1995. pp. 185–89. In Chinese.

46. Islam MT, Croll D, Gladieux P, Soanes DM, Persoons A, Bhattacharjee P, et al. Emergence of wheat blast in Bangladesh was caused by a South American lineage of *Magnaporthe oryzae*. BMC Biology. 2016; 14(1):84, doi: 10.1186/s12915-016-0309-7

47. Steele K, Humphreys E, Wellings C, Dickinson M. Support for a stepwise mutation model for pathogen evolution in Australasian *Puccinia striiformis* f. sp. *tritici* by use of molecular markers. Plant Pathology. 2001; 50(2):174–180.

48. Hovmøller M, Justetson A. Rates of evolution of avirulence phenotypes and DNA markers in a northwest European population of *Puccinia striiformis* f. sp. *tritici*.Molecular Ecology. 2007; 16(21):4637–4647.

49. Jiménez-Gasco M, Milgroom M, Jiménez-Díaz R. Stepwise evolution of races in *Fusarium oxysporum* f. sp. *ciceris* inferred from fingerprinting with repetitive DNA sequences. Phytopathology. 2004; 94(3):228–235.

50. Wang C, Guncar G, Forwood J, Teh T, Catanzariti A, Lawrence GJ, et al. Crystal structures of flax rust avirulence proteins AvrL567-A and -D reveal details of the structural basis for flax disease resistance specificity. Plant Cell. 2007; 19(9):2898–2912.

51. Jia Y, Valent B, Lee FN. Determination of host responses to *Magnaporthe grisea* on detached rice leaves using a spot inoculation method. Plant Disease. 2003; 87(2):129–133.

52. Tai T, Tanksley SD. A rapid and inexpensive method for isolation of total DNA from dehydrated plant tissue. Plant Molecular Biology Reporter. 1990; 8(4):297–303.

53. Rozas J, Sánchez-Del BJ, Messeguer X, Rozas R. DnaSP, DNA polymorphism analyses by the coalescent and other methods. Bioinformatics. 2003; 19(18):2496–2497.

54. Clement M, Posada D, Crandall K. TCS: a computer program to estimate gene genealogies. Molecular Ecology. 2000; 9(10):1657–1659.

55. Fontaine C, Lovett PN, Sanou H, Maley J, Bouvet J-M. Genetic diversity of the shea tree (Vitellaria paradoxa C.F. Gaertn), detected by RAPD and chloroplast microsatellite markers. Heredity. 2004; 93(6):639–648

